# Horizontal Gene Transfers Underpin Ribose Heterotrophy and Central Carbon Metabolism Remodeling in Gloeobacteraceae

**DOI:** 10.1101/2025.11.10.686926

**Authors:** Edi Sudianto, Denis Baurain, Luc Cornet

## Abstract

Gloeobacterales has long been considered a “living fossil” cyanobacterial order, owing to its lack of thylakoid membranes and basal phylogenetic position. However, our study reveals that Gloeobacterales actively integrate horizontally transferred genes into their core metabolism. In Gloeobacteraceae—one of the two families within the order—these genes encode a complete ribose ATP synthase binding cassette (ABC) importer and downstream enzymes, enabling the heterotrophic uptake of external ribose and its assimilation into central carbon metabolism, along with photosynthesis, indicative of photomixotrophy. Beyond ribose utilization, their central carbon metabolism exhibits a mosaic architecture shaped by the integration of foreign genes into the Calvin-Benson-Bassham cycle, the pentose phosphate pathway, and the Embden-Meyerhof-Parnas pathway. Uniquely, these genes appear to have been acquired through multiple independent transfer events, as reflected by their dispersed genomic locations and diverse bacterial donors, including other cyanobacteria and Pseudomonadota. These findings contradict the long-standing view of Gloeobacterales as metabolically primitive relics. Instead, Gloeobacterales is likely a dynamic lineage that continues to adapt and evolve through metabolic innovation and the assimilation of foreign genes into its genomes.

## Introduction

Cyanobacteria are the only prokaryotes capable of oxygenic photosynthesis. This process relies on a linear electron transfer chain to convert light energy and CO_2_ into carbohydrates and release O_2_ as a byproduct (Johnson 2016). Photosynthesis is estimated to have originated ∼3.4 billion years ago in the last common ancestor of the cyanobacteria (Fournier et al. 2021). Owing to this ancient innovation, cyanobacteria are traditionally regarded as strictly photosynthetic organisms (Smith et al. 1967). However, both experimental and environmental evidence have increasingly demonstrated that photomixotrophy—the ability to perform photosynthesis while assimilating additional carbon sources—is more widespread than previously assumed within the phylum (Muñoz-Marín et al. 2024). The first large-scale study by Rippka et al. (1979) in the 1970s found that diverse cyanobacteria are capable of photomixotrophy on a range of carbon sources, including glucose, fructose, sucrose, ribose, and glycerol. For instance, *Nostoc punctiforme* PCC 73102 can metabolize ribose, glucose, fructose, and sucrose (Rippka et al. 1979; Stebegg et al. 2023). These substrates are typically funneled into central carbon metabolic pathways, such as the pentose phosphate pathway (PPP), the Entner-Doudoroff (ED) pathway, the Embden-Meyerhof-Parnas (EMP) pathway, and the Calvin-Benson-Bassham (CBB) cycle (Muñoz-Marín et al. 2020; Stebegg et al. 2023; Lucius and Hagemann 2024).

In most cyanobacteria, photosynthesis occurs within specialized intracellular membranes known as thylakoids. Members of the earliest-diverging order, Gloeobacterales, lack these membranes and instead localize photosynthetic complexes to the cytoplasmic membrane (Raven and Sánchez-Baracaldo 2021; Cornet 2025). This distinctive feature, together with their basal phylogenetic position, has led to the view that Gloeobacterales represent an ancestral state of cyanobacterial evolution—a “living fossil” lineage preserving early photosynthetic traits (Cornet 2025). Despite their ecological diversity, Gloeobacterales remain poorly characterized due to their recalcitrance to cultivation; only four strains have been successfully isolated since their discovery in the 1980s (Grettenberger 2021; Raven and Sánchez-Baracaldo 2021; Saw et al. 2021). Here, we report the first genomic evidence of photomixotrophy within the Gloeobacteraceae, expanding the known metabolic repertoire of this basal cyanobacterial lineage. We further show that the central carbon metabolism of the Gloeobacterales has been shaped by horizontal gene transfer events that likely occurred after the diversification of Gloeobacteraceae and Anthocerotibacteraceae. These findings challenge the notion of Gloeobacterales as a strictly ancestral cyanobacterial lineage, highlighting the dynamic evolutionary history underlying their metabolic networks.

## Results

### Phylogeny and Hierarchical Orthologous Group data summary

Out of the 251 single-copy genes (SCGs) in the cyanobacterial reference profile Hidden Markov Models (pHMMs) of GToTree, our taxa contained 98**–**250 of the SCGs (Table S1). Non-photosynthetic taxa possessed fewer SCGs than the photosynthetic ones (Table S1); specifically, “*Candidatus* Margulisbacteria” (104**–**142), “*Candidatus* Sericytochromatia” (112**–**149), and Vampirovibrionophyceae (98**–**150) contained fewer SCGs than photosynthetic cyanobacteria (159–250). The phylogeny reconstructed from these SCGs showed clear delineation of the major cyanobacterial orders, with strong bootstrap support, particularly among basal lineages (Figure S1). Using the proteomes of the 343 selected taxa, we reconstructed and inferred 41,823 Hierarchical Orthologous Groups (HOGs) at the root node (N0 in Figure S1). For downstream analyses, we identified 34 HOGs of interest corresponding to 37 selected genes primarily involved in ribose uptake and metabolism and central carbon metabolism (Table S2).

### Ribose uptake in Gloeobacteraceae

Our HOG analysis revealed that all Gloeobacteraceae taxa included in this study possess three genes associated with an ATP-synthase Binding Cassette (ABC) transporter system. These three genes are clustered with genes annotated as fructose importers (*frtA*, *frtB*, and *frtC*) in other cyanobacteria (see Table S3) and code for an ATP-binding protein, a substrate-binding protein, and a permease protein, respectively. Interestingly, no Anthocerotibacteraceae taxa have any genes being clustered in these three HOGs.

Based on structural alignment analysis (Table S4) comparing the substrate-binding protein to reference proteins, we determined that this protein is likely a ribose importer (TM-score: 0.94; Root Mean Square Deviation (RMSD): 1.5) rather than a fructose importer (TM-score: 0.75–0.77; RMSD: 2.09**–**2.38). This interpretation is also supported by KEGG (based on sequence homology and the presence of specific motifs), which annotates the three genes as ribose transporter components (e.g., *rbsA* in *G. violaceus*: https://www.kegg.jp/entry/gvi:glr0808).

These results indicate that all examined Gloeobacteraceae taxa possess the complete genetic repertoire required to assemble a functional ribose importer protein. Thus, the seven Gloeobacteraceae taxa included in this study are likely capable of utilizing ribose as an external carbon source in addition to performing oxygenic photosynthesis. In contrast, the absence of these genes in the Anthocerotibacteraceae suggests that the ribose heterotrophy is restricted to the Gloeobacteraceae and was potentially acquired only after it diverged from its sister family (Figure 1). This discovery provides the first genomic evidence of heterotrophic capability in the basal cyanobacteria and demonstrates that photomixotrophy in Gloeobacteraceae is a derived metabolic innovation rather than an ancestral trait.

**Figure 1.**
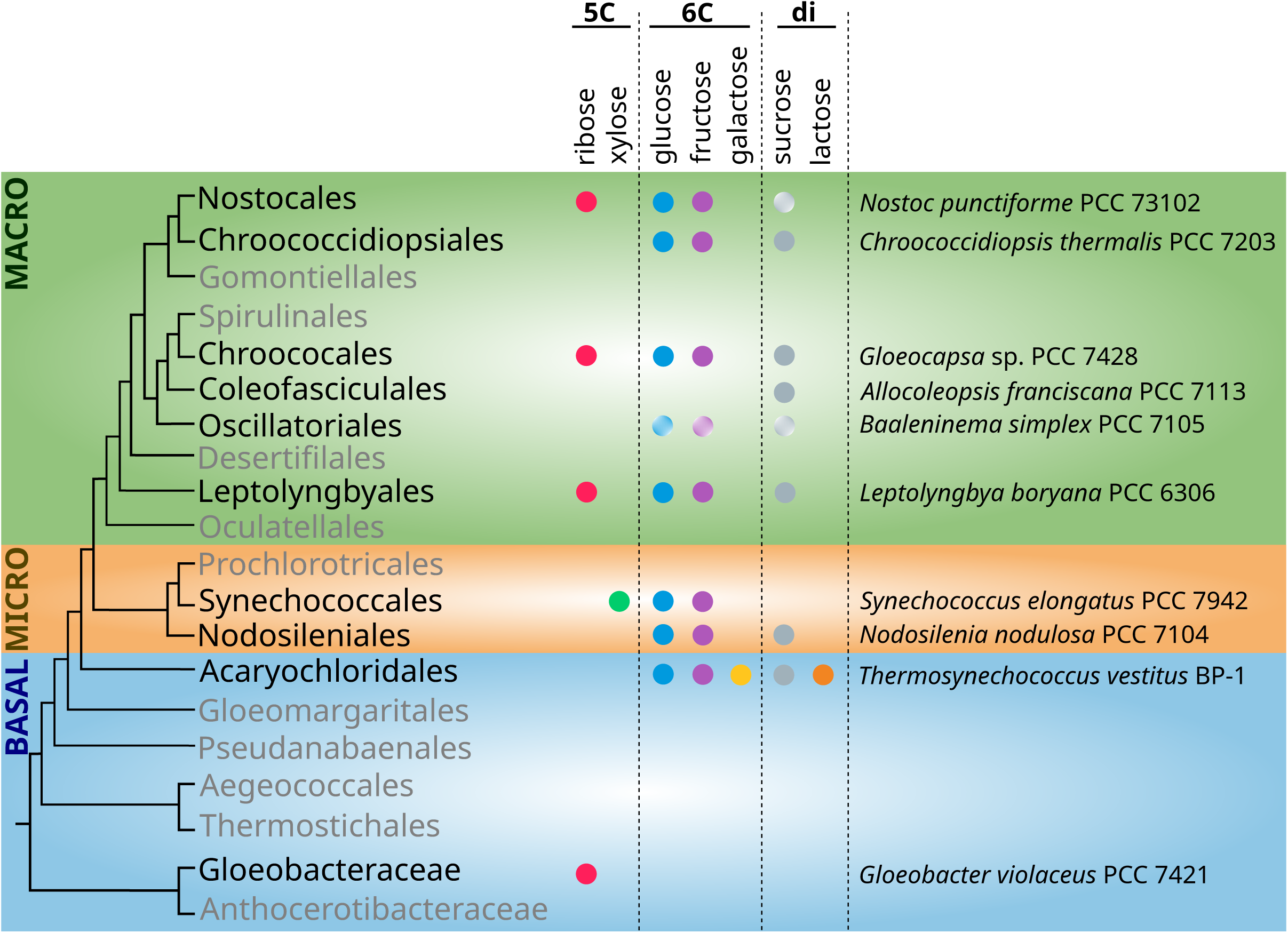
Overview of heterotrophic capability among 19 cyanobacterial orders. The listed taxa and their heterotrophic capability (except for ribose heterotrophy in Gloeobacteraceae) were derived from Stebegg et al. (2023). The phylogeny backbone follows the classification of Strunecký et al. (2023). The Gloeobacterales order is divided into two families, Gloeobacteraceae and Anthocerotibacteraceae, to highlight differences in heterotrophic capability. Orders without known heterotrophic taxa are marked as gray. Circles with a white gradient indicate “weak” ability to utilize the listed sugars. 5C: five carbon monosaccharides (pentoses); 6C: six carbon monosaccharides (hexoses); di: disaccharides. BASAL: basal cyanobacteria, MICRO: microcyanobacteria, MACRO: macrocyanobacteria, following Gisriel et al. (2023).

### Gene gain and loss mosaics in the central carbon metabolism of Gloeobacterales

We found clear signatures of gene gain and loss events across the central carbon metabolism (Figure 2). The three genes responsible for the ribose importer are likely products of horizontal gene transfer (HGT) into an ancestor of Gloeobacteraceae, as they are absent from the HOGs of Anthocerotibacteraceae and other basal cyanobacteria (Table S3). Similarly, the ribokinase (*rbsK*) and ribose-5-phosphate isomerase B (*rpiB*) HOGs are exclusive to the *Gloeobacter* genus and Gloeobacteraceae, respectively, suggesting independent HGT events (Table S3). Although all Gloeobacterales taxa possess a single *rpiA* homolog, phylogenetic analysis indicates that the Gloeobacteraceae copy is non-cyanobacterial (Figure 3).

**Figure 2.**
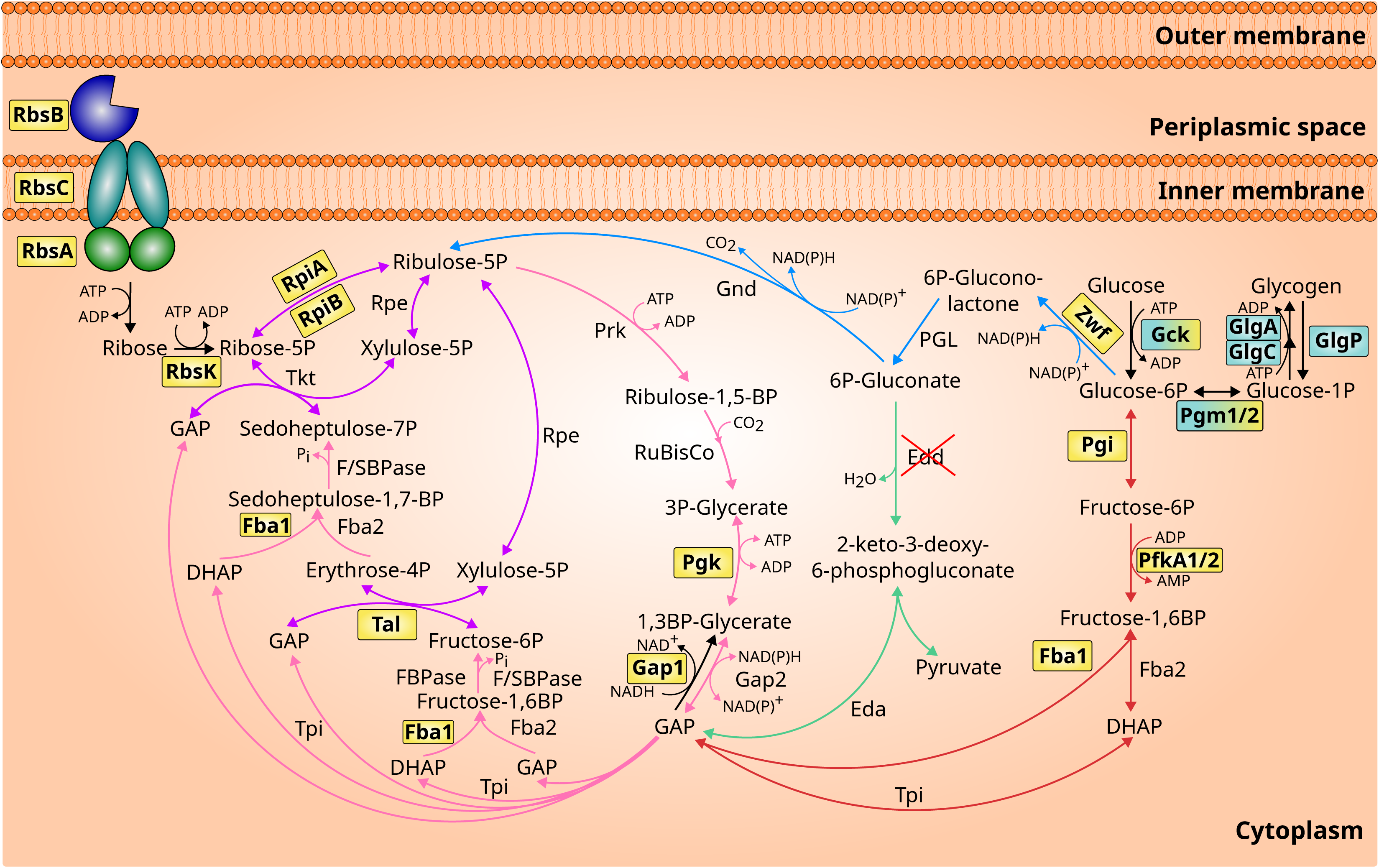
Summary of the central carbon metabolism pathways involved in ribose import and utilization in Gloeobacterales. Arrow colors denote distinct pathways: the Calvin-Benson-Bassham (CBB) cycle is *pink*, the pentose phosphate pathway (PPP) is *blue*, shared pathways between CBB and PPP is *purple*, the Entner-Doudoroff pathway (ED) is *green*, and the Embden–Meyerhof–Parnas pathway (EMP) is *red*. Key enzymes potentially acquired through horizontal gene transfer are highlighted in yellow boxes, while enzymes lost in some taxa are highlighted in cyan boxes. Enzymes exhibiting signatures of horizontally acquired genes before being lost in some taxa are shown in yellow–cyan boxes. An enzyme absent in all Gloeobacterales taxa is marked with a red cross. Illustration based on metabolic pathway information from Bowyer and Leegood (1997) and Caliebe et al. (2025). RbsA: ribose import ATP-binding protein; RbsB: ribose import substrate-binding protein; RbsC: ribose import permease protein; RbsK: ribokinase; RpiA: ribose-5-phosphate isomerase A; RpiB: ribose-5-phosphate isomerase B; Rpe: ribose-5-phosphate epimerase; Prk: phosphoribulokinase; RuBisCo: ribulose-1,5-bis-p carboxylase/oxygenase; Pgk: phosphoglycerate kinase; Gap: glyceraldehyde 3-phosphate dehydrogenase; Tpi: triose-phosphate isomerase; Fba: fructose-bisphosphate aldolase; FBPase: fructose-1,6-bisphosphatase; F/SBPase: Bifunctional fructose-1,6/sedoheptulose-1,7-bisphosphatase; Tal: transaldolase; Tkt: transketolase; Gnd: 6-phosphogluconate dehydrogenase; Pgl: 6-phosphogluconolactonase; Zwf: glucose-6-phosphate 1-dehydrogenase; Edd: 6-phosphogluconate dehydratase; Eda: 2-keto-3-deoxygluconate-6-phosphate (KDPG) aldolase; Pgi: glucose-6-phosphate isomerase; PfkA: phosphofructokinase A; Gck: Glucokinase; Pgm: phosphoglucomutase; GlgA: glycogen synthetase; GlgC: glucose-1-phosphate adenylyltransferase catalytic site; GlgP: glycogen phosphorylase.

**Figure 3.**
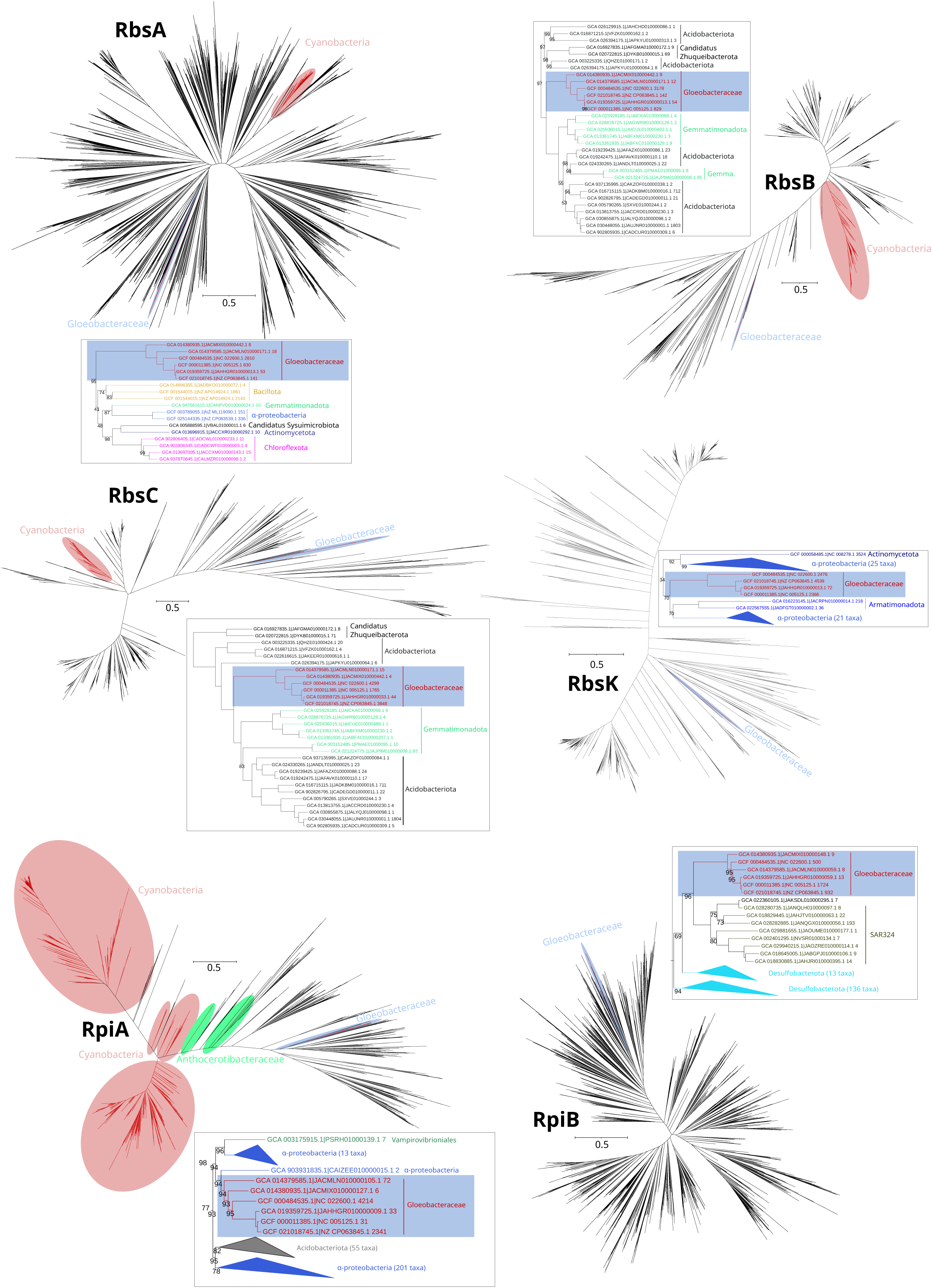
Gene trees of six genes involved in ribose metabolism. The red, blue, and green ellipses show the position of cyanobacteria, Gloeobacteraceae, and Anthocerotibacteraceae taxa, respectively. The tree snippets beside each gene tree display the potential donor of horizontal gene transfer (HGT) to Gloeobacteraceae. Only nodes with bootstrap supports of <100% are shown. These maximum likelihood trees were reconstructed using LG+F+G substitution models and 1,000 ultrafast bootstrap replicates in the following alignment dimensions: RbsA (3,738 sequences ✕ 1,329 sites); RbsB (1,942 sequences × 4,108 sites); RbsC (1,797 sequences × 2,895 sites); RbsK (784 sequences × 398 sites); RpiA (1,834 sequences × 434 sites); RpiB (3,740 sequences × 386 sites). RbsA: ribose import ATP-binding protein; RbsB: ribose import substrate-binding protein; RbsC: ribose import permease protein; RbsK: ribokinase; RpiA: ribose-5-phosphate isomerase A; RpiB: ribose-5-phosphate isomerase B.

In Gloeobacterales, additional HGT-derived genes were identified across the CBB cycle, the PPP, and the EMP pathways, including *pgk*, *gap1*, *fba1*, *tal*, *zwf*, *pgi*, and *pfkA* (Figure S2). In contrast, several genes (*gck*, *pgm1/2*, *glgA*, *glgC*, and *glgP*) are absent in subsets of Anthocerotibacteraceae (Figure 2; Table S3). HGT analysis (Figure S2) indicates that *gck* and *pgm1/2* show signs of horizontally acquired genes before being lost in some members of Anthocerotibacteraceae. Nevertheless, our HOG data show that the majority of cyanobacterial taxa, including Gloeobacteraceae, still retain these genes (Table S3). The *edd* gene was absent from all cyanobacterial genomes, while *pfkA* was detected only in three Anthocerotibacteraceae genomes, potentially indicating a recent acquisition event. Overall, our findings suggest that the central carbon metabolism in Gloeobacterales has been shaped by HGT and differential gene loss, resulting in a mosaic metabolic architecture.

### Ribose metabolism genes in Gloeobacteraceae are mostly acquired via horizontal gene transfer

Phylogenetic analyses of the six ribose import and metabolism genes (including *rbsA, rbsB, rbsC, rbsK, rpiA,* and *rpiB*) demonstrate that all Gloeobacteraceae sequences were acquired through HGT (Figure 3). Anthocerotibacteraceae lack these ribose-related genes entirely, except for *rpiA* (Figure 3).

Surprisingly, the inferred donors are diverse, with only *rbsB* and *rbsC* likely originating from the same source, possibly Acidobacteriota or Gemmatimonadota. The gene trees indeed show that *rbsB* and *rbsC* share the same donor lineage, suggesting co-transfer as a part of a gene cluster. *RbsA,* however, likely has a distinct evolutionary origin. There is no clear definition of the *rbsA* donor as its gene tree places the Gloeobacteraceae sequences as sister to a group composed of diverse bacteria, including Bacillota, Gemmatimonadota, Alphaproteobacteria, “*Ca.* Sysuimicrobiota”, Actinomycetota, and Chlorofexota. Thus, it is predicted that the donor of *rbsA* is distinct from those of *rbsB* and *rbsC*.

Furthermore, our gene trees clearly indicate Alphaproteobacteria as the donor of *rbsK* and *rpiA* and SAR324/Desulfobacterota as the donor of *rpiB* in Gloeobacteraceae. *RpiA* represents an especially interesting case, where the HGT copy appears to have replaced the endogenous cyanobacterial copy still present in Anthocerotibacteraceae (Figure 3). Together, these findings strongly imply that all genes involved in ribose import and metabolism in Gloeobacteraceae were independently acquired via HGT from multiple bacterial sources, underscoring the evolutionary plasticity of this basal cyanobacterial lineage.

### Contrasting the genetic repertoire of Gloeobacteraceae and Anthocerotibacteraceae

To elucidate the evolutionary dynamics of genes involved in ribose import and metabolism and central carbon metabolic pathways across Gloeobacterales, we inferred patterns of gene acquisition and loss, along with their putative donor lineages (Figure 4). This analysis revealed marked contrast in the evolutionary trajectories of Gloeobacteraceae and Anthocerotibacteraceae.

**Figure 4.**
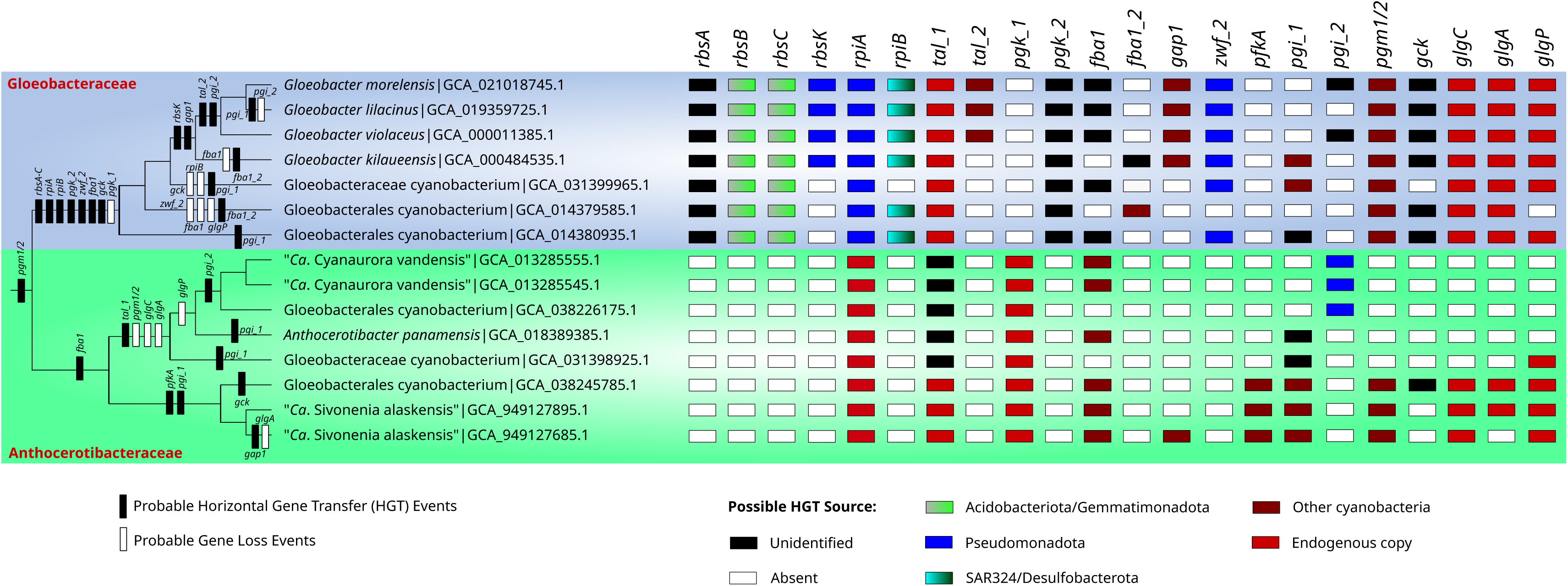
Proposed model of horizontal gene transfer (HGT) and gene loss events for genes involved in central carbon metabolism among the two Gloeobacterales families, as indicated in Figure 2. The Gloeobacterales phylogeny is based on the cyanobacterial phylogeny in Figure S1. Colored boxes on the right denote the gene presence/absence and their potential HGT donor. It is noteworthy that three genomes are named either Gloeobacterales cyanobacterium (GCA_038226175.1, GCA_038245785.1) or Gloeobacteraceae cyanobacterium (GCA_031398925.1) while they clearly belong to Anthocerotibacteraceae in our species tree (Figure S1).

In Gloeobacteraceae, numerous horizontally transferred genes were inferred to have been acquired at the base of the family. These include *rbsA–C, rpiAB, pgk_2, zwf_2,* and *fba1,* accompanied by the loss of *pgk_1*. These genes likely originated from phylogenetically diverse donors (Figure 4; Figure S2), such as Acidobacteriota/Gemmatimonadota (*rbsBC*), Pseudomonadota (*rpiA* and *zwf_2*), and SAR324/Desulfobacterota (*rpiB*). The donors of *rbsA, pgk_2,* and *fba1* remain unidentified. Subsequent acquisitions include *rbsK* (putatively from Pseudomonadota) and *gap1* (other cyanobacteria) in the common ancestor of the four *Gloeobacter* species. Additional transfers, such as *tal_2* (cyanobacterial origin) and *pgi_2* (unknown origin), occurred later within the *G. violaceus*–*G. morelensis* lineage.

In contrast, the Anthocerotibacteraceae is characterized by a single inferred HGT event (*tal_1*) and multiple gene losses (*pgm1/2, glgC,* and *glgA*) in the clade containing “*Ca*. Cyanaurora vandensis” and *Anthocerotibacter panamensis.* These taxa appear to have replaced their native transaldolase (*tal_1*) gene with an unclassified HGT-derived copy. Additional losses include *glgP,* which is absent from four species in the family but retained in one Gloeobacterales genome (GCA_031398925.1). Three taxa, including two “*Ca.* Cyanaurora vandensis” and a Gloeobacterales cyanobacterium (GCA_038226175.1), acquired *pgi_2* from the Pseudomonadota. Note that despite being annotated as Gloeobacterales cyanobacterium or Gloeobacteraceae cyanobacterium (GCA_038226175.1, GCA_038245785.1, and GCA_031398925.1), these genomes are confidently placed within Anthocerotibacteraceae according to our species tree (Figure S1). Furthermore, the “*Ca.* Sivonenia alaskensis” clade uniquely carries a *pfkA* gene of other cyanobacteria origin, while this gene is absent from all other Gloeobacterales taxa examined. These patterns suggest that Gloeobacteraceae underwent frequent horizontal gene acquisition and metabolic innovation, whereas Anthocerotibacteraceae followed a conservative trajectory with selective gene loss and limited HGT, reflecting divergent trends of metabolic evolution within the order.

### Frequent gene rearrangements and genome plasticity in Gloeobacterales

Most horizontally transferred genes in Gloeobacterales are dispersed rather than organized in operons or conserved clusters. Synteny comparisons among the four complete genome assemblies of Gloeobacterales, including *G. violaceus, G. morelensis, G. kilaueensis,* and *Anthocerotibacter panamensis,* revealed limited conservation of local gene neighborhoods (Figure 5).

**Figure 5.**
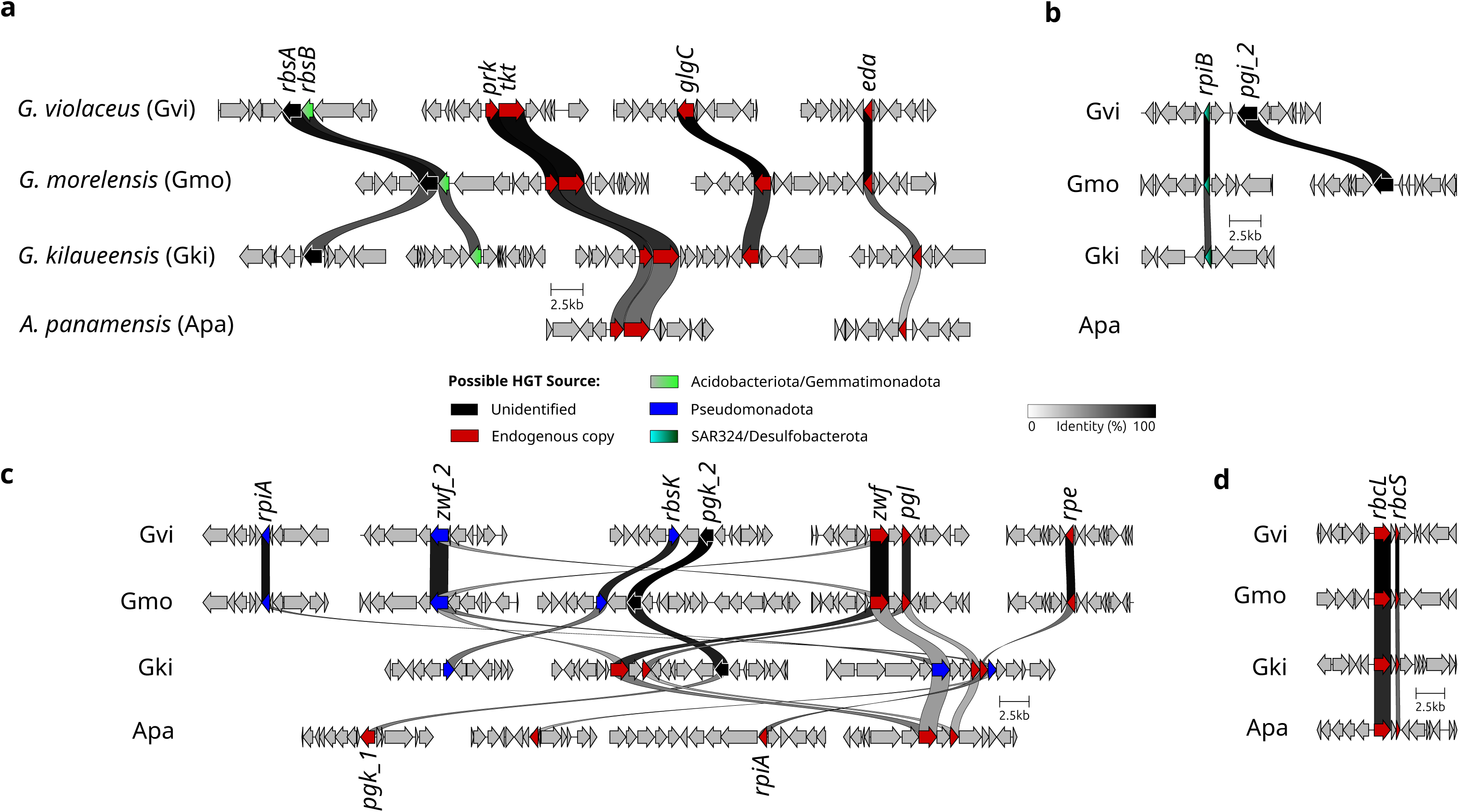
Synteny of selected central carbon metabolism genes across the four complete genomes of Gloeobacterales that are currently available. The illustrated regions contain both endogenous (red) or horizontally transferred genes (color corresponding to donor lineage), including: (a) *rbsA-rbsB*, (b) *rpiB-pgi_2*, (c) *rpiA, rbsK, zwf_2*, and (d) the endogenous *rbcL-rbcS*. Only regions with two or more neighboring genes in at least one of the four genomes are shown. Gvi: *Gloeobacter violaceus*; Gmo: *Gloeobacter morelensis*; Gki: *Gloeobacter kilaueensis*; Apa: *Anthocerotibacter panamensis*.

The ribose importer genes *rbsA* and *rbsB* were adjacent only in *G. violaceus* and *G. morelensis*, but not in *G. kilaueensis*. Even native cyanobacterial gene arrangements, such as *prk-tkt, glgC,* and *eda,* differed markedly among the *Gloeobacter* genomes (Figure 5a). Similarly, the *pgi_2* gene was located near the *rpiB* gene in *G. violaceus* but not in *G. morelensis* (Figure 5b).

Genes acquired from the same lineages also displayed variable genomic positions. For example, *rpiA* and *zwf_2,* both originating from Pseudomonadota, are located close to one another in *G. kilaueensis* but are situated in distant genomic regions in *G. violaceus* and *G. morelensis* (Figure 5c). In contrast, some ancestral operons, such as the *rbcL-rbcS,* have retained synteny across the family (Figure 5d). Collectively, these results suggest that horizontally transferred genes are integrated in the Gloeobacterales genomes and subject to frequent rearrangements, reflecting ongoing post-acquisition reshuffling rather than stable incorporation into conserved metabolic loci.

## Discussion

### Ribose is an important carbon source for Gloeobacteraceae

Previous studies (White and Shilo 1975; Rippka et al. 1979; Stebegg et al. 2023) have identified some cyanobacteria capable of heterotrophic ribose utilization, including members of the Nostocales, Chroococales, Leptolyngbyales, and Geitlerinematales (Figure 1). Here, we provide the first genomic evidence of a similar capability in the basal cyanobacterial lineage Gloeobacteraceae, inferred from the presence of ribose ABC importer genes in all examined genomes (Table S3). This finding suggests that the capacity to import and metabolize ribose emerged after the divergence from Anthocerotibacteraceae but before the diversification of Gloeobacteraceae (ca. <∼1.4 billion years ago–>300 million years ago; Rahmatpour et al. (2021)). Phylogenetic analyses (Figure 3) indicate that these genes were likely horizontally transferred from at least two distinct donors: (1) either Acidobacteriota/Gemmatimonadota for the ATP-binding and permease components, and (2) unresolved bacterial taxa for the ribose-binding protein. The non-contiguous arrangement of ribose importer genes in Gloeobacteraceae (Figure 5) further supports multiple, independent acquisition events. Such a chimeric ABC transporter is unusual, as they are typically transferred as an intact unit, e.g., maltose transporters in Thermococcales and Thermotogales (Noll et al. 2008) and phosphate transporters in Firmicutes (Moreno-Letelier et al. 2011). The gene fragmentation pattern, however, is consistent with the extensive genome rearrangements typical of cyanobacteria, which often lack conserved synteny even in stable operons such as the *dcw* cluster (Léonard et al. 2022), secondary metabolite genes (Entfellner et al. 2022), and EPS synthesis genes (Kim et al. 2024).

Ribose is among the earliest and most abundant sugars in nature (Schönheit et al. 2016). In its phosphorylated form (ribose-5P, R5P), it plays central roles in nucleotide, amino acids, DNA, and RNA synthesis (Schönheit et al. 2016; Tang et al. 2021), the central carbon metabolism (e.g., CBB cycle and PPP; Figure 2; Noor et al. (2010), Makowka et al. (2020), Lucius and Hagemann (2024)), and antibiotic resistance (Seregina et al. 2024; Seregina et al. 2025). A ribose ABC importer is present in diverse bacteria, including *Escherichia coli* (Clifton et al. 2015)*, Bacillus subtilis* (Woodson and Devine 1994), and *Salmonella typhimurium* (Aksamit and Koshland 1972), underscoring the broad importance of ribose uptake and metabolism.

The functional implications of this ribose uptake system are particularly intriguing in light of the respiratory characteristics of *G. violaceus*. Koyama et al. (2008) reported exceptionally high respiratory activity—ca. twentyfold higher than *Synechocystis* PCC 6803—suggesting significant reliance on respiration. However, the substrate driving this oxygen consumption has remained enigmatic (Koyama et al. 2008). Our findings offer a possible explanation for this pattern. The presence of a complete ribose uptake and catabolic system within the Gloeobacteraceae suggests a capacity for photomixotrophy, in which ribose could serve as an auxiliary carbon source. Imported ribose may subsequently be oxidized via the PPP to generate reducing power for respiration, thereby sustaining energy metabolism when oxygenic photosynthetic electron flow is limited. This interpretation is consistent with the elevated oxygen uptake and may help explain the slow but sustained growth of *G. violaceus* in culture.

The presence of both *rpiA* and *rpiB* genes in Gloeobacteraceae further highlights the importance of ribose metabolism. These enzymes catalyze the interconversion between R5P and ribulose-5P (Ru-5P), a key metabolic step linking ribose utilization with the central carbon network. Both proteins share largely the same function but are structurally distinct (Chen et al. 2020). The *rpiA* gene is nearly universal among organisms, but *rpiB* occurs mainly in Actinobacteria and diverse eukaryotes (Tang et al. 2021; Santana-Molina et al. 2025). Only a few bacteria, such as *E. coli* and *Listeria monocytogenes*, encode both forms (Maltsev et al. 2005; Chen et al. 2020; Tang et al. 2021). It was proposed that while *rpiA* primarily catalyzes the conversion of Ru-5P to R5P, the reverse conversion is preferentially performed by *rpiB* under high R5P concentrations (Tang et al. 2021). Thus, the presence of *rpiB* in Gloeobacteraceae is likely linked to elevated intracellular R5P levels resulting from active ribose import.

Collectively, the presence of all key genes required for ribose import (*rbsA–C*) and its assimilation (*rbsK*, *rpiA,* and *rpiB*) into the central carbon metabolism suggests that heterotrophic ribose utilization is an integral metabolic feature of Gloeobacteraceae. These genes were likely acquired through multiple independent HGT events rather than a single operon transfer. The discovery that ribose heterotrophy may fuel the high respiratory activity of *G. violaceus* provides a physiological basis for this genomic adaptation. These findings reveal that Gloeobacteraceae, long viewed as strictly photoautotrophic “living fossils”, have in fact evolved a more versatile and dynamic carbon metabolism through HGT, combining an ancient photosynthetic machinery with a more recently acquired heterotrophic capability.

### Horizontal gene transfer and gene loss reshape central carbon metabolism in Gloeobacterales

Shi and Falkowski (2008) suggested that the “core” genes of cyanobacteria (such as those involved in photosynthesis and the CBB cycle) are largely conserved; most of the HGT acquisition affects the “shell” or accessory genes. However, several studies have reported HGT in the “core” central carbon metabolism in some cyanobacteria, including the acquisition of *rbcL-rbcS* from ɑ-proteobacteria in marine Synechococcaceae (Cabello-Yeves et al. 2022), *fba1* from red algae in marine Synechococcaceae (Rogers et al. 2007; Godde et al. 2018), and *gap1* from *Anabaena* in *G. violaceus* (Figge et al. 1999). Our analyses further expand the list of potential HGTs in the central carbon metabolism of Gloeobacterales, identifying *tal, pgk, fba1, pgi, pfkA*, and *zwf* as additional candidates (Figure 2; Figure 4). The lack of *edd* in Gloeobacterales verified findings by Evans et al. (2024) that suggested the absence of a complete ED pathway in cyanobacteria and plants.

Phylogenetic reconstructions (Figure S2) confirm that *gap1* was horizontally transferred from other cyanobacteria in all four examined *Gloeobacter* species, consistent with Figge et al. (1999). Additional transfers from cyanobacterial donors are clear for *tal_2, fba1, fba1_2, pfkA*, *pgi_1,* and *pgm1/2* (Figure 4). These transfers are relatively “recent”, postdating the diversification of donor lineages. This finding underscores other cyanobacteria as major contributors of metabolic genes to Gloeobacterales. Members of Pseudomonadota also appear as frequent donors, contributing *rbsK, rpiA,* a second copy of *zwf* (in Gloeobacteraceae), and *pgi_2* (in Anthocerotibacteraceae). Similar Pseudomonadota-to-cyanobacteria transfers have been previously reported, such as RuBisCo–carboxysome operon from ɑ-proteobacteria (Cabello-Yeves et al. 2022) and nitrogen fixation operon from δ-proteobacteria (Chen et al. 2022). Conversely, even with extensive searches across more than 100,000 representative genomes in GTDB, some HGT donors remained unidentifiable, possibly due to limited resolution or incomplete taxonomic representation in the current database.

The ecological context of these transfers is consistent with the lifestyle of Gloeobacterales. Cyanobacteria often form dense biofilms in microbial mats (Bhaya et al. 2025; Bozan et al. 2025), which promote DNA exchanges through close physical contact with other bacteria. Indeed, Gloeobacterales microbial mats typically harbor various cyanobacterial taxa, including Nostocales, Leptolyngbyales, Synechococcales, and Chroococcales (Cuzman et al. 2010; Pessi et al. 2023), as well as abundant Pseudomonadota (Pessi et al. 2023). The dominance of these two phyla within Gloeobacterales-associated mats parallels their role as principal HGT donors. Other putative donors, such as Acidobacteriota and Gemmatimonadota, are also commonly found in microbial mats containing Gloeobacterales taxa (Zeng et al. 2020; Powell et al. 2024).

The absence of glycogen metabolism genes in certain members of the Anthocerotibacteraceae (Figure 4) represents an intriguing metabolic feature. Glycogen serves as the primary storage molecule for photosynthetically fixed carbon in cyanobacteria, thereby buffering against fluctuations in energy and carbon (Xu et al. 2013; Shinde et al. 2020). Interestingly, the lack of glycogen metabolism has been frequently reported in parasitic, symbiotic, or fastidious bacteria (Henrissat et al. 2002). Mutants of *glgC* and *glgA* in *Synechocystis* and *Synechococcus* exhibit severe defects in glycogen accumulation (Xu et al. 2013; Carrieri et al. 2017). The loss of these genes leads to several physiological consequences, including: (1) a prolonged lag phase in initiating photosynthesis following dark periods (Shinde et al. 2020), (2) excretion of soluble sugars or “energy spilling” (Gründel et al. 2012; Xu et al. 2013), and (3) increased susceptibility to environmental stress, particularly under fluctuating dark– light or photomixotrophic conditions (Gründel et al. 2012). Alternatively, members of Anthocerotibacteraceae lacking canonical *glg* genes may possess alternative carbon storage routes, such as those mediated by Rv3032-, GlgE-, or glucosylglycerol-based pathways observed in other bacteria (Chandra et al. 2011; Ortega-Martínez et al. 2025).

These findings demonstrate that HGT and gene loss have been major drivers of metabolic reorganization in Gloeobacterales. The repeated recruitment of cyanobacterial and proteobacterial genes suggests that Gloeobacterales did not simply conserve ancestral metabolic traits but instead actively restructured their physiology through foreign gene assimilation.

### Evolutionary and ecological drivers of metabolic innovation in Gloeobacterales

*Gloeobacter* is remarkable among cyanobacteria as it respires more than it photosynthesizes, indicating a strong dependence on external organic carbon sources (Koyama et al. 2008). The absence of thylakoid membranes inherently limits their photosynthetic efficiency (Rahmatpour et al. 2021), which explains both their slow growth and the difficulty of cultivating them in the laboratory. When the photosynthetic electron transport chain first emerged in the cytoplasmic membrane of ancestral organisms—possibly resembling modern *Gloeobacter*—this innovation would have conferred a significant evolutionary advantage (Cornet 2025). However, as thylakoid-bearing cyanobacteria evolved and diversified within the same microbial mats (Cuzman et al. 2010; Pessi et al. 2023), Gloeobacterales may have faced increasing ecological competition, and the lack of thylakoid membranes became a liability over time.

The evidence presented here suggests that Gloeobacteraceae subsequently evolved toward photomixotrophy, secondarily acquiring the capacity to exploit exogenous organic carbon sources such as ribose, to compensate for the lack of competitiveness from other cyanobacteria. This transition was facilitated by multiple horizontal gene transfers, producing a patchwork of imported and rearranged genes even within core metabolic pathways. The genomic mosaicism observed in their central metabolism, spanning sugar importers, the CBB cycle enzymes, and the PPP components, underscores the evolutionary plasticity of these basal cyanobacteria. More importantly, these reorganizations challenge the perception of Gloeobacterales as “primitive” cyanobacteria merely preserving ancestral traits. Instead, they represent an ancient lineage that has adapted to metabolic and ecological pressures through gene exchange and functional innovation.

## Conclusion

Our findings show that HGT has profoundly shaped the evolution of Gloeobacteraceae, enabling them to persist in low-light, nutrient-variable habitats by coupling inefficient photosynthesis with heterotrophic ribose utilization. HGT has also remodeled the Gloeobacterales’ central carbon metabolism by combining endogenous genes with horizontally-acquired genes, resulting in mosaic metabolic pathways. The microbial mats inhabited by Gloeobacterales likely serve as active gene-exchange networks, with other cyanobacteria and Pseudomonadota acting as the primary donors. This reinterpretation positions Gloeobacterales not as a static relic of early cyanobacterial evolution, but as metabolically versatile survivors whose genomes record a long history of innovation; thus, the ancestral nature of Gloeobacterales should be interpreted with caution.

## Materials and Methods

### Taxa selection and phylogenetic tree reconstruction

We curated a dataset comprising 343 cyanobacteria and related non-photosynthetic taxa encompassing the full diversity of cyanobacterial orders described by Strunecký et al. (2023). In brief, we downloaded the cyanobacteria genome list from GTDB release R220 (Parks et al. 2022) and selected one to two representatives for each genus, prioritizing those with a high CheckM (Parks et al. 2015) completeness score (>80%) and a low contamination level (<10%). To better represent the basal cyanobacteria, we included all available genomes from NCBI for the orders Gloeobacterales, Thermostichales, Pseudanabaenales, and Gloeomargaritales. The Pseudanabaenales dataset was further dereplicated using ToRQuEMaDA v0.2.1 (Léonard et al. 2021) with the following parameters: “p=200, l=200, t=0.80, j=12, a=100, r=10, d=1, min=1, type=taxonomic, alg=JI, egn=jellyfish”. Additionally, the taxon sampling was enriched with strains reported to exhibit heterotrophic capacity. In total, our data comprises 104 non-photosynthetic taxa (including “*Candidatus* Margulisbacteria”, “*Ca.* Sericytochromatia”, and Vampirovibrionophyceae) and 239 (photosynthetic) cyanobacteria (Table S1).

We used GToTree v1.8.8 (Lee 2019) and the cyanobacterial SCG pHMMs available in the tool to reconstruct the SCG supermatrix based on 251 individual alignments for the 343 selected taxa. The pipeline uses MUSCLE v5.1 (Edgar 2022) and TrimAl v.1.4.rev15 (Capella-Gutiérrez et al. 2009) for sequence alignment and trimming, respectively. We slightly modified the parameters of GToTree by adding the following options: “-c 1 and -G 0”. The supermatrix resulting from concatenated SCG alignments was used to infer a phylogenomic tree using IQ-TREE v2.3.6 (Minh et al. 2020) with the automatic best substitution model (-m MFP; Kalyaanamoorthy et al. 2017) and 1,000 ultrafast bootstrap (Hoang et al. 2018) replicates. The final tree is shown in Figure S1.

### Hierarchical Orthologous Group inference and ortholog search

The 343 selected genomes were first annotated using the Prodigal v2.6.3 (Hyatt et al. 2010) function implemented in the GEN-ERA toolbox v3.0.0 (Cornet et al. 2023). The orthologous gene inference was subsequently performed using OrthoFinder v2.5.4 (Emms and Kelly 2019) together with the previously reconstructed phylogeny (Figure S1) as the guide tree. The inferred HOGs were used to identify orthologs of genes of interest, including those related to heterotrophy and the central carbon metabolic pathways (Table S2). Reference genes were obtained from the literature (e.g., *Nostoc* fructose importer (Ekman et al. 2013) and *Synechocystis* fructose-1,6-biphosphatase (Caliebe et al. 2025)) or the KEGG database (Kanehisa et al. 2023). The presence or absence of each gene across HOGs is summarized in Table S3.

### Ribose structural alignment analyses

We used US-align v20241108 (Zhang et al. 2022) to assess the substrate specificity of the transporter gene identified in this study. We first downloaded the predicted protein structure of *Gloeobacter violaceus*’ substrate-binding transporter (AlphaFold Protein DB: Q7NMF9) from the AlphaFold Protein Structure Database (Varadi et al. 2022). Reference proteins with known substrate specificities, e.g., for fructose, glucose, galactose, ribose, and maltodextrin (Table S4), were also retrieved from either the RSCB Protein Data Bank (Berman et al. 2000) or AlphaFold Protein Structure Database (Varadi et al. 2022). Each reference protein was aligned to the *G. violaceus* substrate-binding protein using US-align. Both the TM-score and RMSD were recorded. A perfectly aligned protein yields a TM-score = 1 (Zhang and Skolnick 2004), while RMSD < 2 Å indicates a high structural similarity (Shamsian et al. 2024).

### Identification of horizontal gene transfer events

We first extracted the cyanobacterial sequences within the 34 identified HOGs of interest in this study (Table S3). Next, these protein sequences were aligned using MAFFT v7.5.2 (Katoh et al. 2002). The resulting alignments were converted into pHMMs using the hmmbuild function in HMMER v3.4 (http://hmmer.org/). These pHMMs were then searched against the GTDB R220 bacteria representative genome dataset (107,235 species) using the hmmsearch function. To reduce redundancy, we filtered the hits using OmpaPa v0.252040 (https://metacpan.org/dist/Bio-MUST-Apps-OmpaPa), retaining approximately 704–5,206 sequences for downstream analyses. These sequences were subsequently realigned using MAFFT and trimmed with Clipkit v2.4.1 in the default (smart-gap) parameter (Steenwyk et al. 2020) before gene tree reconstruction using IQ-TREE v2.3.6. We set the IQ-TREE parameters to 1,000 ultrafast bootstrap replicates and the LG+F+G substitution model (for computational efficiency). The resulting trees were color-formatted using the format-tree.pl function in Bio-MUST-Core v0.252040 (https://metacpan.org/dist/Bio-MUST-Core) and visualized using iTOL v7 (Letunic and Bork 2024).

Because the initial HMM profiles were constructed from all cyanobacterial sequences within each HOG, some putative HGTs into Gloeobacterales may have been masked. To mitigate this effect and improve donor identification, we repeated the above workflow with a modification: the pHMMs were constructed using only the Gloeobacterales sequences available in the HOGs. The remaining steps, such as GTDB search, OmpaPa filtering, and gene tree reconstruction, followed the same workflow described above.

### Synteny analyses

To investigate the genomic context of the horizontally transferred genes, we examined the synteny of those HGT-acquired genes using clinker v0.0.31 (Gilchrist and Chooi 2021). Four Gloeobacterales taxa with complete genome assemblies were selected for comparison, including *G. violaceus, G. morelensis, G. kilaueensis,* and *Anthocerotibacter panamensis*. For each taxon, genomic regions encompassing the gene of interest (typically ±5 kb flanking regions) were extracted using an in-house script (https://doi.org/10.6084/m9.figshare.30422521). These regions were compared and visualized using clinker, which identifies homologous genes based on translated sequence similarity and aligns genomic loci accordingly (Gilchrist and Chooi 2021).

## Acknowledgements

We thank Raphaël Léonard for the help in running the ToRQuEMaDA software and Louise Hambücken for assistance in identifying key primary literature relevant to this study.

## Author Contributions

ES: Data curation; formal analysis; investigation; visualization; writing – original draft; writing – review & editing. DB: Conceptualization; funding acquisition; software; writing – review & editing. LC: Conceptualization; funding acquisition; supervision; writing – review & editing.

## Data Availability

All genomic data, generated data, and custom scripts used in this study are deposited in the FigShare repository (https://doi.org/10.6084/m9.figshare.30422521) under the CC BY 4.0 license. The GTDB R220 bacteria representative genome dataset can be downloaded from the GTDB website (https://gtdb.ecogenomic.org/downloads).

