## Supplementary figures and images for "Horizontal Gene Transfers Underpin Ribose Heterotrophy and Central Carbon Metabolism Remodeling in Gloeobacteraceae"

### Supplemental Figures

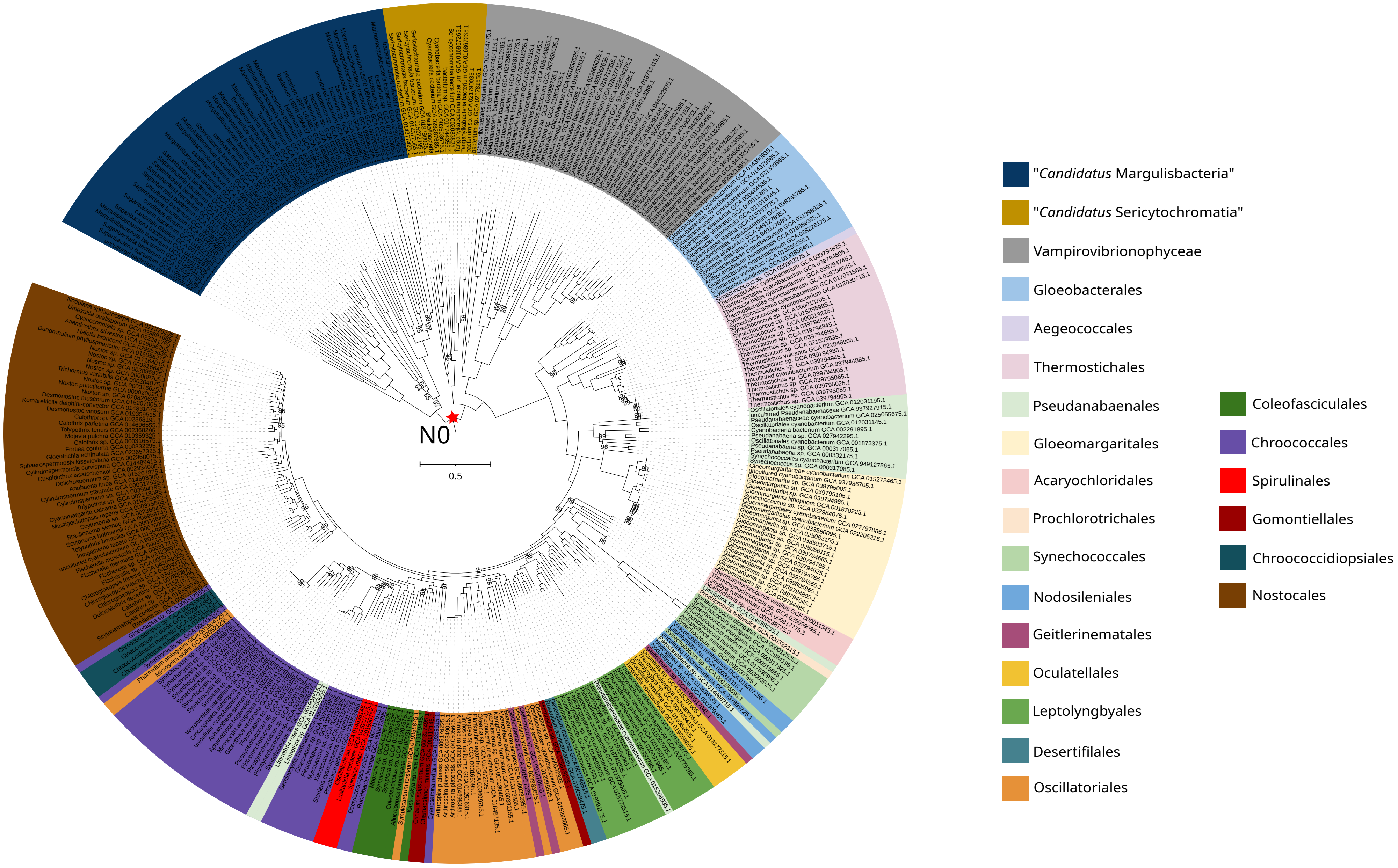

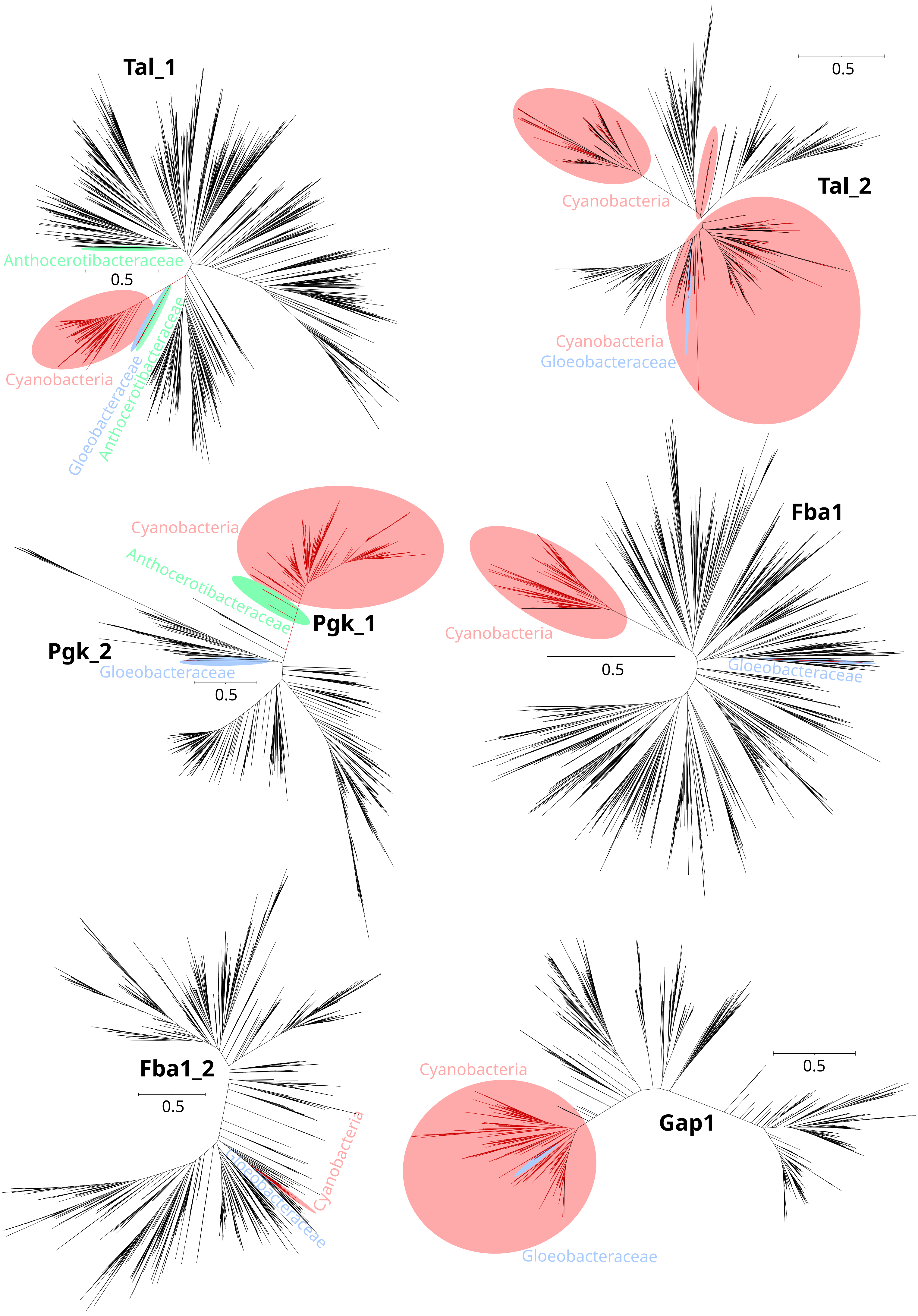

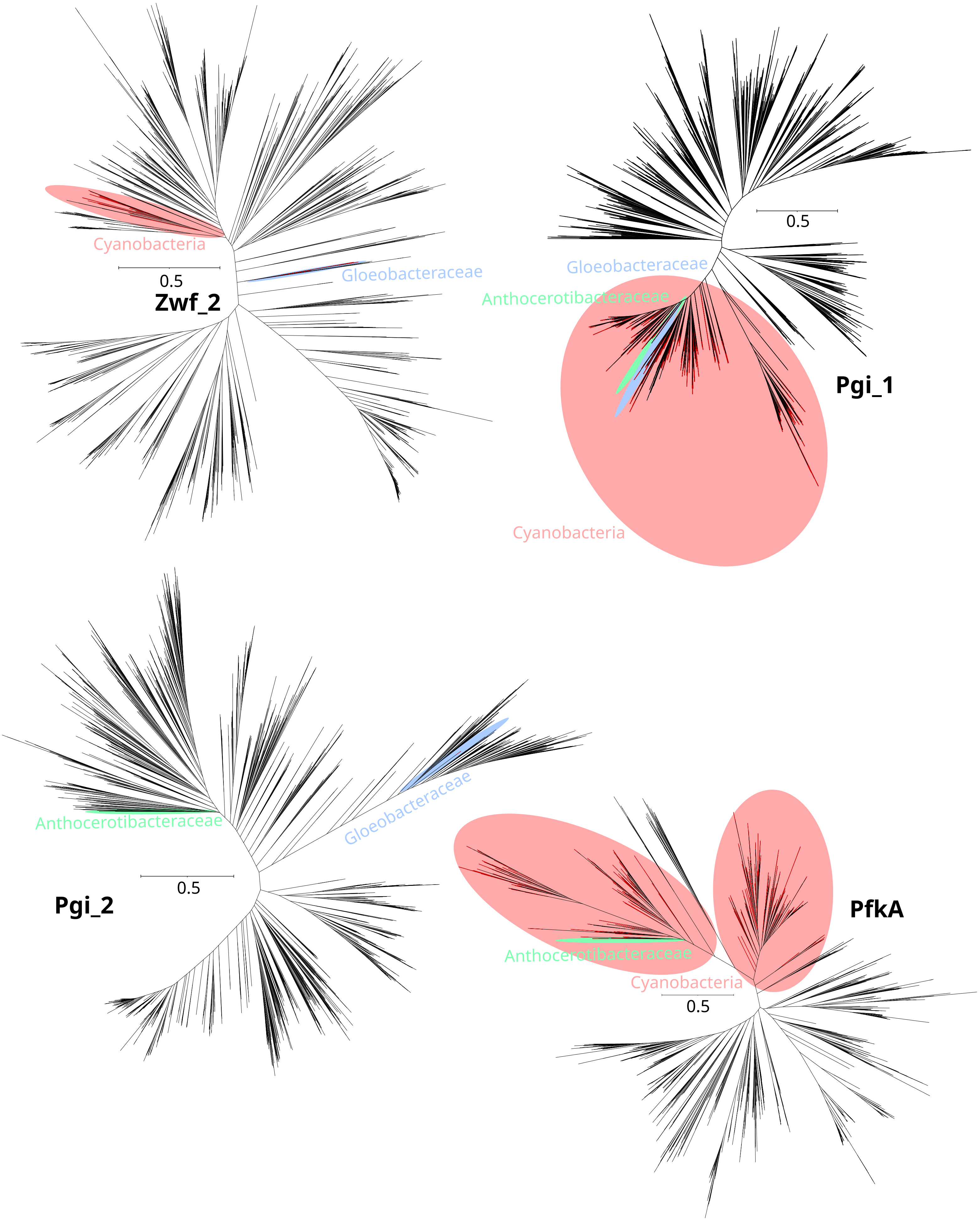

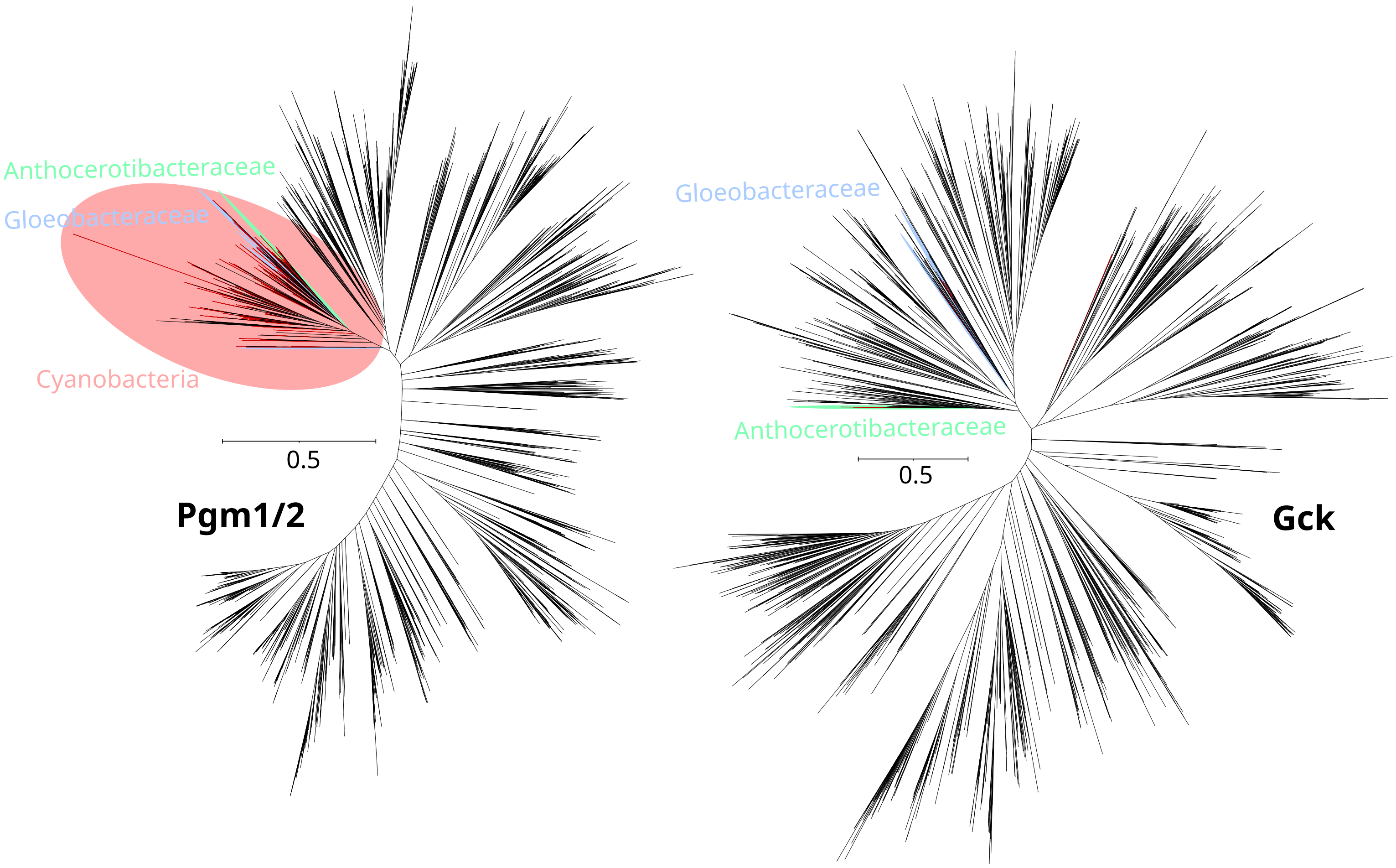
